# Immunostimulatory effects of *Bacillus coagulans* SANK70258

**DOI:** 10.1101/2023.02.21.529342

**Authors:** Yuki Ikeda, Naoto Ito, Niya Yamashita, Natsuki Minamikawa, Kazuki Nagata, Takuya Yashiro, Masanori Aida, Ryouichi Yamada, Masakazu Hachisu, Chiharu Nishiyama

## Abstract

Specific intestinal bacteria modulate immunoresponses through various pathways, and several probiotic bacteria have been identified as immunostimulants by screening. In the present study, we evaluated the immunomodulating effects of *Bacillus coagulans* SANK70258 (*B. coagulans* SANK70258), a spore-forming and lactic acid-producing bacterium usable as food supplement for human and animals. We found that treatment of mouse splenocytes with γ-ray irradiated *B. coagulans* SANK70258 induced high amount of IFN-γ in comparison with 7 kinds of typical lactic acid bacteria. Further analyses using splenocytes revealed that NK cell is a major source of IFN-γ, and *B*. *coagulans* SANK70258-induced IFN-γ production was inhibited by neutralization of IL-12 or IL-23, depletion of CD11c^+^ cells, and inhibition of NFκB. *B. coagulans* SANK70258 also induced release of IFN-γ from activated CD8^+^ T cells, and increased expression of chemokine receptors in CD8^+^ T cells. *B. coagulans* SANK70258-treatment induced production of cytokines from bone marrow-derived dendritic cells, which is reduced by knockdown of *Tlr2* and *Nod2. B. coagulans* SANK70258-treatment also induced IgA production from Peyer’s patch cells with high level among tested lactic bacteria. The oral intake of γ-ray irradiated *B. coagulans* SANK70258 significantly increased intestinal IgA levels and IgA-expressing B cells in the Peyer’s patch of mice. Taken together, we conclude that *B. coagulans* SANK70258 possesses high activity as immunostimulant inducing production of IFN-γ and IgA.

## Introduction

The intestinal immune homeostasis is in a balance of fight against infectious pathogens and tolerance toward commensal microbiota and food ingredients. Specific intestinal bacteria modulate immunoresponses through various pathways, such as activation of innate immune cells by binding of bacterial cell components to receptors recognizing pathogen-associated molecular patterns, and production of the secondary metabolites including short-chain fatty acids (SCFAs) that regulate gene expression and function of immune-related cells.

*Bacillus coagulans* is a bacteria species, whose intake alleviates the pathology of intestinal diseases, such as constipation and colitis, and has recently attracted the attention with its usefulness in food processing industry due to the resistance to heat temperature and acidic condition depending on the spore-forming character [1]. *Bacillus coagulans* SANK70258 (*B. coagulans* SANK70258), also termed *Weizmannia coagulans* SANK70258, is a lactic acid bacterium used as probiotics for livestock animals and humans. Oral administration of *B. coagulans* SANK70258 effectively promotes the growth of broiler chickens with conferring the protection ability against *Coccidia* infection [2, 3] and modulates the composition of SCFAs in the intestine [4]. A study using a model culture system of human colonic microbiota revealed that *B*.

*coagulans* SANK70258 increased the concentration of butyrate and number of *Lachnospiraceae* bacteria in the intestine and reduced colonic *Enterobacteriaceae* species [5]. Although these findings support the beneficial effects of *B. coagulans* SANK70258 on the host body, the roles of *B. coagulans* SANK70258 on the host immune systems are still unclear.

In the current study, we investigated the roles of *B. coagulans* SANK70258 as an immunostimulant by using *in vitro* and *in vivo* experiments, and revealed that *B. coagulans* SANK70258 cell components exhibit high activity in production of IFN-γ and IgA from immune-related cells.

## Materials and Methods

### Mice and Cells

The present study was approved by the Animal Care and Use Committees of Tokyo University of Science (K22005, K21004, K20005, and K19006), and was conducted in accordance with the guideline of the Institutional Review Board of Tokyo University of Science. The spleen, the Peyer’s patch, and the bone marrow were isolated from Balb/c mice (Japan SLC, Hamamatsu, Japan) to obtain splenocytes, Peyer’s patch cells, and to generate BMDCs, respectively. BMDCs were developed from BM cells as previously described [6]. To neutralize cytokines, anti-IL-12p40 (clone C17.8, BioLegend), anti-IL-23p19 (clone MMp19B2, BioLegend), anti-IL-6 (clone MP5-20F3, BioLegend), and isotype control (#RTK2071, BioLegend) Abs were used. Depletion of CD11c^+^ cells from splenocytes was performed by autoMACS Pro Separator (Miltenyi Biotec) with CD11c MicroBeads UltraPure mouse (Miltenyi Biotec). CD8^+^ T cells were isolated from the spleen using Mojosort Magnetic Separation System CD8 Naïve T cell Isolation Kit (BioLegend). Anti-CD3ε Ab (clone 145-2111C, BioLegend) and anti-CD28 Ab (clone 37.51, TONBO Bioscience) were used to stimulate CD8^+^ T cells. BAY11-7082, LE540 (#123-04521, Wako), Brefeldin A (#420601, BioLegend) were used as inhibitors.

### Preparation of lactic acid bacteria

*B. coagulans* SANK70258 and 7 kinds of bacteria (*Lactobacillus antri* 15950, *L. sakei subsp. sakei* 1157, *L. buchneri* 1068, *L. gasseri* 1131, *L. plantarum subsp. Plantarum* 1149, *Leuconostoc mesenteroides subsp. cremoris* 16943, *L. mesenteroides subsp. cremoris* 6124; all obtained from RIKEN BRC) were cultured in MRS medium for 1 day under a same shaking condition. After washing with saline, harvested bacteria were freeze-dried and were γ-ray irradiated.

### ELISA

The concentrations of mouse IFN-γ, IL-12p40, IL-6, and IL-10, were determined by ELISA using ELISA MAX Deluxe Sets (BioLegend), and IgA concentration was measured by mouse IgA uncoated ELISA kit (Invitrogen) or mouse IgA ELISA Quantification Set (#E90-103, Bethyl Laboratories).

### Flow cytometry

To identify CD4^+^ T cells, CD8^+^ T cells, and NK cells in whole splenocytes, cells were stained with anti-mouse CD3ε-PerCP (clone 145-2C11, BioLegend), anti-mouse CD4-FITC (clone GK1.5, BioLegend), anti-CD8-VioGreen (clone 53-6.7, Miltenyi Biotec), anti-NK1.1-PE (clone PK136, BioLegend), and anti-CD49b-APC (clone DX5, BioLegend) Abs. Intracellular IFN-γ was stained with anti-mouse IFN-γ-PE/Cyanine7 (clone XMG1.2, BioLegend) after treatment with Fixation buffer (#420801, BioLegend) and Intracellular staining perm wash buffer (#421002, BioLegend). A MACS Quant analyzer (Miltenyi Biotec) and Flowjo (Tomy Digital Biology, Tokyo, Japan) were used to detect fluorescence and to analyze data, respectively.

### Small interfering RNA

Electroporation was performed to introduce siRNA into BMDCs using a mouse dendritic cell nucleofector kit (Lonza, Basel, Switzerland) and a Nucleofector 2b (Lonza). Following siRNAs were purchased from Invitrogen (Carlsbad, CA, USA): *Tlr2* (#MSS216272), *Tlr4* (#MSS211922), *Nod2* (#MSS217440), Stealth RNAi siRNA Negative Control Lo GC (#12935-200) as for negative control of *Tlr4* siRNA, Stealth RNAi siRNA Negative Control Hi GC (#12935-400) for *Tlr2* and *Nod2* siRNAs.

### Quantification of mRNA

Complementary DNA was synthesized from total RNA, which was isolated from BMDCs using the ReliaPrep RNA Cell Miniprep System (#Z76012, Promega, Madison, USA), using ReverTra Ace qPCR Master Mix (#FSQ-201, TOYOBO, Osaka, Japan). Quantitative real-time PCR was performed with Thunderbird SYBR qPCR Mix (#QPS-201, TOYOBO) on the StepOne Real-Time PCR System (Applied Biosystems, Kanagawa, Japan). The nucleotide sequences (from 5’ to 3’) of the primer sets used in qPCR are listed as follows.

*Gzmb*-F; TGCATTCCCCACCCAGACTA, *Gzmb*-R; TTCAGCTTTAGCAGCATGATGTC, *Prf1*-F; GAGTGTCGCATGTACAGTTTTCG, *Prf1*-R; GCGCCTTTTTGAAGTCAAGGT, *Ccr4*-F; GCAAGGCAGCTCAACTGTTCT, *Ccr4*-R; TGGCATTCATCTTTGGAATCG, *Ccr5*-F; GGCTCTTGCAGGATGGATTTT, *Ccr5*-R; GGTGCTGACATACCATAATCGATGT, *Ccr6*-F; TCTGAATGAATTCCACAGAGTCCTACT, *Ccr6*-R; CCATGGTCTGGAGGAATAGAATAATAC, *Ccr9*-F; GCACTTCCCCTCCTGAAGCT, *Ccr9*-R; CTTGTGAGTTCTGTGGGGCATCA, *Cxcr3*-F; TGCCAAAGGCAGAGAAGCA, *Cxcr3*-R; CATCTAGCACTTGACGTTCACTAACC, *Cxcr6*-F; GAGCACACTTCACTCTGGAACAA, *Cxcr6*-R; CCATCATCCATGGCATCA, *Il1b*-F; AGTTGACGGACCCCAAAGA, *Il1b*-R; GGACAGCCCAGGTCAAAGG, *Tlr2*-F; GAATTGCATCACCGGTCAGAA, *Tlr2*-R; TCCTCTGAGATTTGACGCTTT, *Tlr4*-F; GCTAAGTGCCGAGTCTGAGTGTAA, *Tlr4*-R; TGCAGCCTTTCAGAAACACATT, *Nod2*-F; CGTGCGCCTGCTCCAT, *Nod2*-R; CACCCTCAGGGACAAGAAGTTC.

Primers for *Il6* [7] and *Aldh1a2* mRNAs [8] were described in our previous reports.

### Statistical analysis

A two-tailed Student’s t-test was used to compare two samples and a one-way ANOVA followed the Tukey-Kramer multiple comparison test was employed to compare multiple (more than three) samples.

## Results and Discussion

### B. coagulans *SANK70258 induces IFN-γ production from NK cells*

To evaluate the effects of *B. coagulans* SANK70258 as immunostimulant, we incubated the mouse splenocytes in the presence of SANK70258 or various lactic acid bacteria, which were sterilized by γ-ray treatment. Determination of IFN-γ concentrations in the culture media of splenocytes after 48 h incubation showed that the SANK70258 component exhibited the highest IFN-γ production ability among tested bacteria components (Fig. 1A), in a dose-dependent manner (Fig. 1B). Then, to identify the IFN-γ-producing cells in whole splenocytes, we performed flow cytometric analysis using Abs against IFN-γ and cell type markers. As shown in Fig. 1C, the SANK70258-treatment markedly increased the frequency of IFN-γ-producing cells in NK population, whereas none and little increase was observed in CD4^+^ T cells and CD8^+^ T cells, respectively.

**Figure 1.**
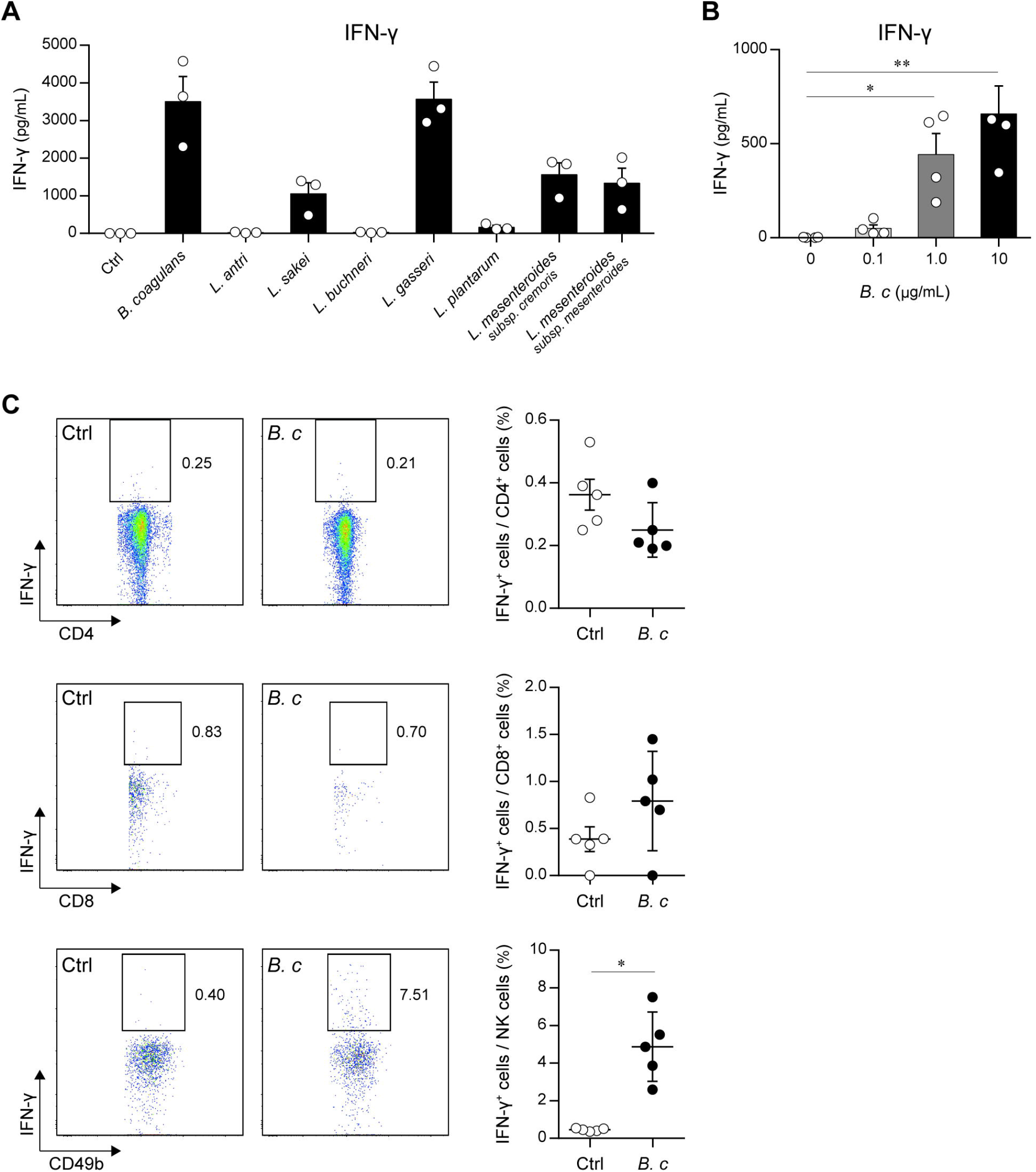
IFN-γ induction activity of lactic acid bacteria and identification of IFN-γ producing cells in splenocytes stimulated with *B. coagulans* SANK70258. **A**. and **B**. Concentrations of IFN-γ in culture media of splenocytes. Whole cells prepared from the spleen of mouse (5 × 10^5^ cells/500 μL) were incubated in the presence or absence of 10 μg/mL γ-ray irradiated bacteria (**A**) or were incubated with indicated amount of *B. coagulans* SANK70258 (**B**) for 48 h. IFN-γ concentrations of culture media were determined by ELISA. The data represent the mean ± SE of 3 independent experiments performed with triplicate samples (**A** and **B**). **C**. Frequencies of IFN-γ producing cells. Whole splenocytes (5 × 10^5^ cells/500 μL) cultured with or without 10 μg/mL *B. coagulans* SANK70258 for 12 h were incubated for additional 5 h in the presence of Monensin and Brefeldin A and were then stained with Abs against IFN-γ and cell-type specific markers. The data represent the mean ± SE of 5 independent experiments. Statistical analysis was performed with two-tailed Student’s *t*-test. \**p* < 0.05.

These results indicate that *B. coagulans* SANK70258 effectively induced IFN-γ release from NK cells mainly.

#### *Participant cells and molecules in* B. coagulans *SANK70258-induced IFN-γ production*

To reveal the molecular mechanisms by which *B. coagulans* SANK70258-treatment induced IFN-γ production, we examined the effects of cytokine blocking, cells depletion, and intracellular-signaling inhibition. As shown in Fig. 2A, addition of anti-IL-12p40 Ab into culture media of spleen cells completely inhibited *B. coagulans* SANK70258-induced production of IFN-γ, and anti-IL-23p19 Ab also significantly suppressed the IFN-γ release. Then, to confirm the involvement of IL-12/IL-23-producing cells in the *B. coagulans* SANK70258-induced IFN-γ production, we compared IFN-γ levels between the whole spleen cells and CD11c^+^-depleted spleen cells, and found that the IFN-γ production following *B. coagulans* SANK70258 treatment was decreased by the depletion of CD11c^+^ cells (Fig. 2B). To further examine whether the NFκB-signaling activated in *B. coagulans* SANK70258-treated cells participate in IFN-γ production, we stimulated whole splenocyte with *B. coagulans* SANK70258 in the presence or absence of BAY11-7082,m an inhibitor of NF-kB. Both production of IFN-γ and IL-12p40 from *B. coagulans* SANK70258-treated splenocyte was completely inhibited by NF-kB inhibition (Fig. 2C). These results demonstrated that *B. coagulans* SANK70258 induced IFN-γ production from NK cells via NF-κB-mediated stimulation of IL-12/23-producible cells including DCs.

**Figure 2.**
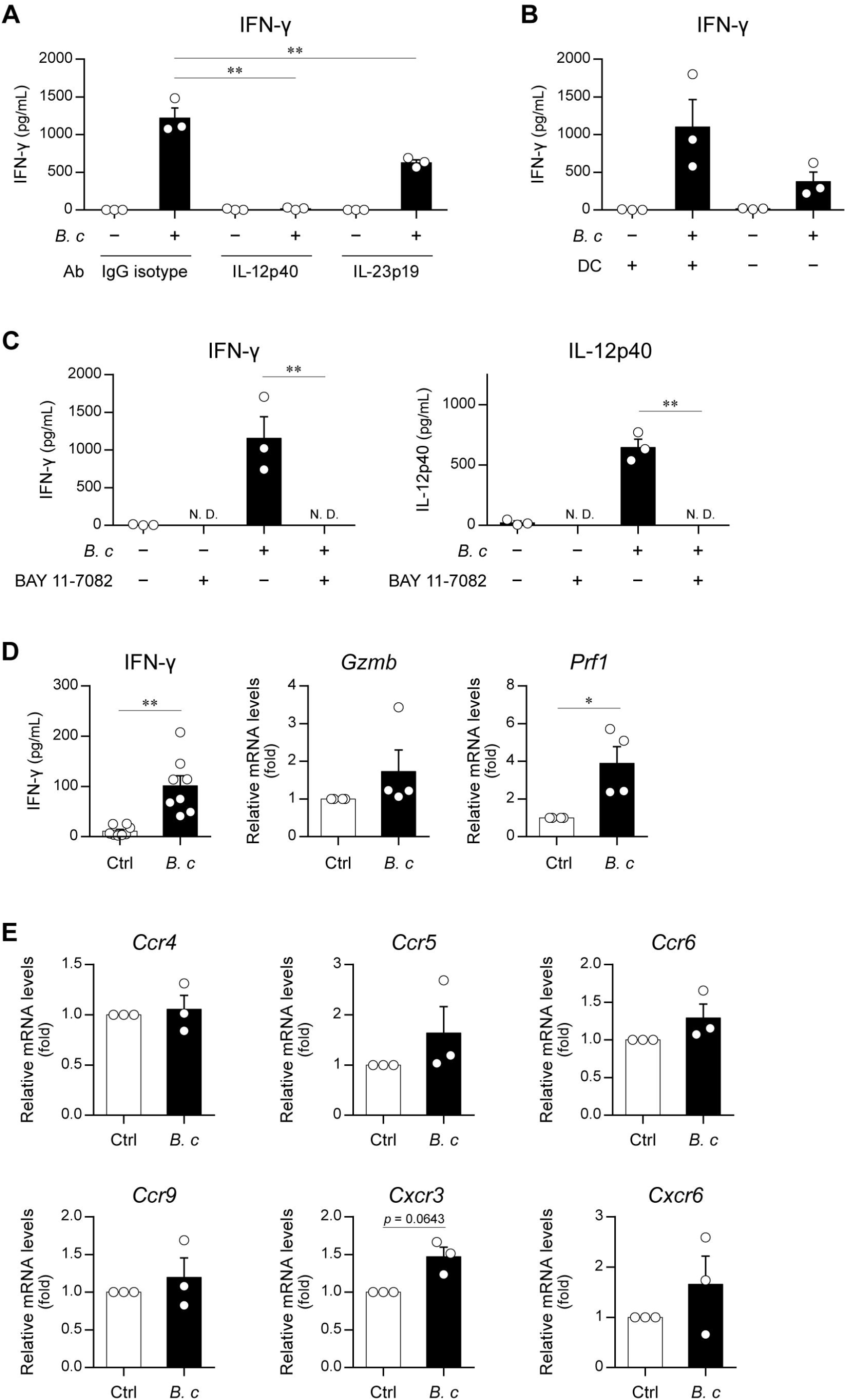
Roles of DCs and CD8^+^ T cells involved in IFN-γ production in *B. coagulans* SANK70258-stimulated splenocytes. **A**. Effects of neutralizing Abs on *B. coagulans* SANK70258-induced IFN-γ production. Splenocytes were incubated with or without 10 μg/mL *B. coagulans* SANK70258 in the presence of 0.5 μg/mL neutralizing Ab or its control for 48 h. **B**. Effects of CD11c^+^ depletion on *B. coagulans* SANK70258-induced IFN-γ production. **C**. Effects of NFκB inhibition on cytokine production in *B. coagulans* SANK70258-treated splenocytes. **D**. *B. coagulans* SANK70258 enhanced IFN-γ production from CD3/CD28-dependently stimulated CD8^+^ T cells. Splenic CD8^+^ T cells were incubated in Ab-coated dishes in the presence or absence of 10 μg/mL *B. coagulans* SANK70258 for 48 h, and IFN-γ concentrations in collected culture supernatants and *Gzmb* and *Prf1* mRNA levels in harvested cells were determined. **E**. Effects of *B. coagulans* treatment on transactivation of chemokine receptor genes in CD8^+^ T cells. Splenic CD8^+^ T cells were cultivated with or without 10 μg/mL *B. coagulans* SANK70258 for 3 h, and then harvested to determine mRNA levels. The data represent the mean ± SD from three independent assays performed in triplicate. *, *p* < 0.05; **, *p* < 0.01. Tukey-Kramer test (**A**-**D**) and Student’s *t*-test (**E** and **F**) were used.

In a cytoplasmic-staining experiment (Fig. 1C), slight expression of IFN-γ was detected in CD8^+^ T cells, which is reported to express TLRs and be able to receive the stimulation by PAMPs [9, 10]. To investigate whether *B. coagulans* SANK70258 directly induces IFN-γ production from CD8^+^ T cells, isolated splenic CD8^+^ T cells were incubated in anti-CD3 Ab- and anti-CD28 Ab-coated dishes in the presence or absence of *B. coagulans* SANK70258. IFN-γ in culture supernatant was significantly increased by *B. coagulans* SANK70258 treatment, and mRNA levels of *Gzmb* and *Prf1* were upregulated in *B. coagulans* SANK70258-treated CD8^+^ T cells (Fig. 2D). In a recent study, CCR6 expression in CD8^+^ T cells was increased by microbial exopolysaccharide produced by *Lactobacillus*, which contributes the antitumor adjuvant effect of the *Lactobacillus* on immune-checkpoint blockade treatment [11]. Then, we quantified mRNA levels of chemokine receptors in *B. coagulans* SANK70258-treated CD8^+^ T cells, and observed that mRNAs of several receptors including CCR6 tended to be increased by *B. coagulans* SANK70258 treatment (Fig. 2E).

#### *Roles of NOD2 and TLR2 in* B. coagulans *SANK70258-dependent cytokine production of DC*

Above-mentioned results using splenocytes indicated that DC is a candidate source of IL-12, which produces IL-12 following *B. coagulans* SANK70258 treatment. To evaluate the effects of *B. coagulans* SANK70258-treatment on DCs, we determined mRNA levels of cytokines in BMDCs incubated with *B. coagulans* SANK70258. Treatment with *B. coagulans* SANK70258 apparently increased mRNA levels of *Il6, Il1b*, and *Aldh1a2* in BMDCs in a dose-dependent manner (Fig. 3A), suggesting that *B. coagulans* SANK70258 directly stimulated DCs. To evaluate the roles of PAMP receptors in *B. coagulans* SANK70258-induced stimulation of DCs, we performed a knockdown experiment using siRNAs for *Tlr2, Tlr4*, and *Nod2*, which are reported to be receptors for lactic acid bacteria components [12]. Although LPS-induced release of cytokines was decreased in *Tlr4* siRNA-transfected BMDCs, knockdown of TLR4 did not reduce cytokine release from *B. coagulans* SANK70258-stimulated BMDCs (Fig. 3B). In contrast, knockdown of TLR2 and NOD2 suppressed the release of IL-6 and IL-12p40 from *B. coagulans* SANK70258-treated BMDCs (Fig. 3C). *B. coagulans* SANK20758 treatment also induced IL-10 release from BMDCs, which is a key character of probiotic bacteria useful for prevention of inflammatory diseases and was markedly reduced by NOD2 knockdown but not by knockdown of TLR2 and TLR4 (Fig. 3B and 3C). Based on the finding that reduced expression of PAMP receptors recognizing peptidoglycan-related components suppressed the cytokine release from *B. coagulans* SANK70258-treated BMDCs.

**Figure 3.**
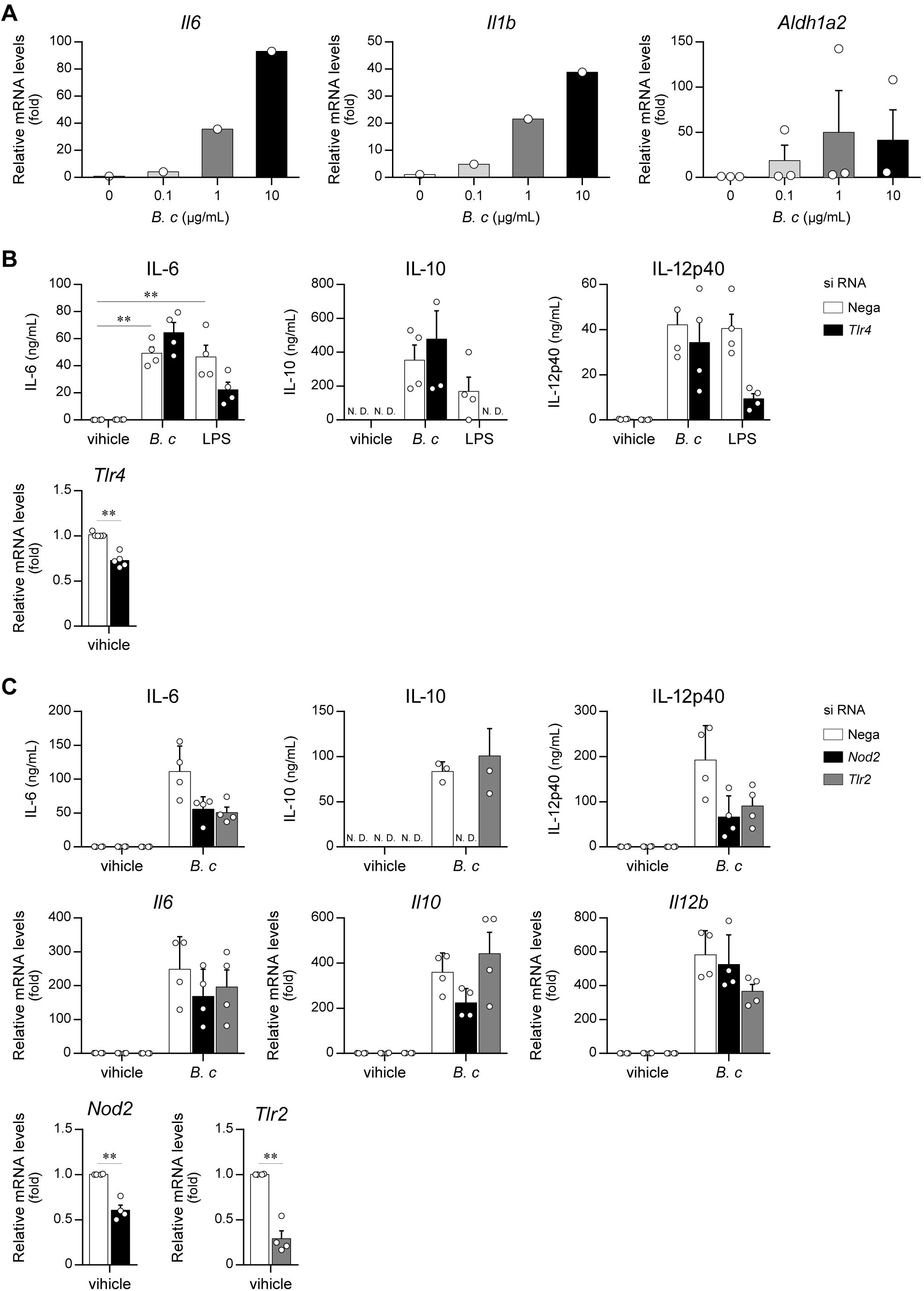
Effects of *B. coagulans* SANK70258 treatment on gene expression in BMDCs. **A**. mRNA levels of *Il6, Il1b*, and *Aldh1a2* in BMDCs. **B**. Effects of *Tlr4* knockdown on *B. coagulans* SANK70258-induced expression of cytokines in BMDCs. **C**. Cytokine production from *Tlr2* or *Nod2* siRNA transfected BMDCs. **D**. Cytokine producing activities of *B. coagulans* SANK70258-derived fractions. To determine mRNA levels in cells and the amounts of cytokines in culture media, cells and culture supernatants were collected 4 h and 24 h after stimulation, respectively. For the stimulation, 10 μg/mL *B. coagulans* SANK70258 or 100 ng/mL LPS was added. BMDCs were cultured for 48 h after siRNA transfection (**B, C**). The data represent the mean ± SD from three independent assays performed in triplicate. *, *p* < 0.05; **, *p* < 0.01. Tukey-Kramer test (**A**-**D**) were used.

### B. coagulans *SANK70258 accelerates IgA production* in vitro *and* in vivo

*B. coagulans* SANK70258 treatment increased the expression of IL-6 and RALDH2 in DCs, which are known to be accelerator of IgA production [13]. To evaluate the effects of *B. coagulans* SANK70258 on IgA production, we cultured whole cells prepared from the Peyer’s patch with *B. coagulans* SANK70258. After 3-7 days cultivation, apparent amounts of IgA accompanied with the production of IL-6 and IFN-γ were detected in *B. coagulans* SANK70258-treated Peyer’s patch cells (Fig. 4A). We also confirmed that *B. coagulans* SANK70258-induced increase of IgA was significantly suppressed in the co-presence of IL-6-neutralizing Ab and an RAR inhibitor (Fig. 4B). When the amounts of IgA in culture media of Peyer’s patch-derived cells incubated with or without lactic bacteria were compared, we found that *B. coagulans* SANK70258 exhibited the highest activity of IgA production among the tested bacteria (Fig. 4C).

**Figure 4.**
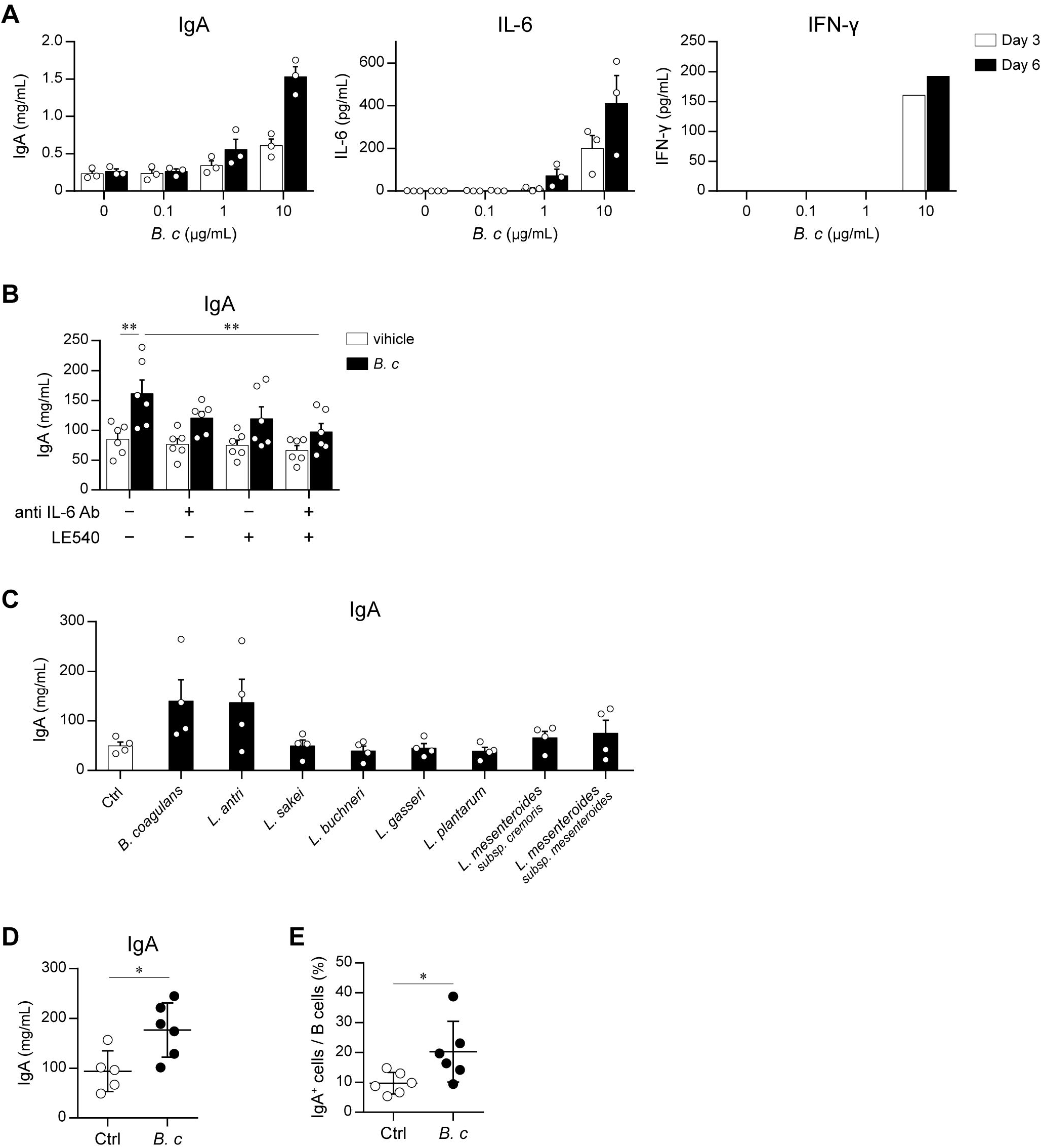
Induction of IgA production by *B. coagulans* treatment *in vitro* and *in vivo*. **A**. The amounts of IgA and cytokine proteins in the Peyer’s patch-derived cells. Whole cells (5 × 10^5^ cells/500 μL) isolated from the Peyer’s patch were incubated with or without indicated concentrations of *B. coagulans* SANK70258 for 3 or 7 days, and concentrations of IgA, IL-6, and IFN-γ in culture supernatant were determined by ELISA. **B**. Effects of IL-6 neutralization and RAR inhibition on IgA production from *B. coagulans* SANK70258-stimulated Peyer’s patch cells. One μg/mL anti-IL-6 Ab and/or 1 μM LE540 were added to culture media of Peyer’s patch-derived cells prior to addition of 10 μg/mL *B. coagulans* SANK70258, and the supernatants after 7 days cultivation were collected to determine IgA concentrations. **C**. IgA concentrations in the culture media of Peyer’s patch cells incubated with or without 10 μg/mL lactic acid bacteria for 7 days. **D**. The amount of IgA proteins in feces. **E**. Frequency of IgA^+^ B cells in the Peyer’s patch. Balb/c mice were fed by 0.2% w/w *B. coagulans* SANK70258-containing or its control diet (**D** and **E**). The data represent the mean ± SEM of individuals (**A**-**E**). *, *p* < 0.05; **, *p* < 0.01. Tukey-Kramer test (**B**) and Student’s *t*-test (**D** and **E**) were used.

Finally, to investigate the effects of *B. coagulans* SANK70258 *in vivo*, we determined the amount of IgA in feces of mice orally administered *B. coagulans* SANK70258. The intake of *B. coagulans* SANK70258 for 2 weeks significantly increased IgA levels in feces (Fig. 4D), and frequency of IgA-producing B cells in the Peyer’s patch (Fig. 4E).

In the present study, we found that *B. coagulans* SANK70258 cell components induce the high amount production of IFN-γ and IgA from immune cells. Although we used γ-ray-treated *B. coagulans* SANK70258 focusing on the roles of cell components as an immunostimulant in the present study, considering that *B. coagulans* SANK70258 can reach the intestine alive by forming spore, it is expected that *B. coagulans* SANK70258 exhibits further beneficial effects on host body via production of the secondary metabolites.

